# Vulnerable Brain Networks Associated with Risk for Alzheimer’s Disease

**DOI:** 10.1101/2022.06.15.496331

**Authors:** Ali Mahzarnia, Jacques A Stout, Robert J Anderson, Hae Sol Moon, Zay Yar Han, Kate Beck, Jeffrey N Browndyke, David B. Dunson, Kim G Johnson, Richard J O’Brien, Alexandra Badea

## Abstract

Brain connectomes provide untapped potential for identifying individuals at risk for Alzheimer’s disease (AD), and can help provide novel targets based on selective circuit vulnerability. Age, APOE4 genotype, and female sex are thought to contribute to the selective vulnerability of brain networks in Alzheimer’s disease, in a manner that differentiates pathological versus normal aging. These brain networks may predict pathology otherwise hard to detect, decades before overt disease manifestation and cognitive decline. Uncovering network based biomarkers at prodromal, asymptomatic stages may offer new windows of opportunity for interventions, either therapeutic or preventive. We used a sample of 72 people across the age span to model the relationship between Alzheimer’s disease risk and vulnerable brain networks. Sparse Canonical Correlation analysis (SCCA) revealed relationships between brain subgraphs and AD risk, with bootstrap based confidence intervals. When constructing a composite AD risk factor based on sex, age, genotype, the highest weight was associated with genotype. Next, we mapped networks associated with auditory, visual, and olfactory memory, and identified networks extending beyond the main nodes known to be involved in these functions. The inclusion of cognitive metrics in a composite risk factor pointed to vulnerable networks, and associated with the specific memory tests. These regions with the highest cumulative degree of connectivity in our studies were the pericalcarine, insula, banks of the superior sulcus and cerebellum. To help scale up our approach, we extended Tensor Network Principal Component Analysis (TNPCA) to evaluate AD risk related subgraphs, introducing CCA components and sparsity. When constructing a composite AD risk factor based on sex, age, and genotype, and family risk factor the most significant risk was associated with age. Our sparse regression based predictive models revealed vulnerable networks associated with known risk factors. The prediction error was 17% for genotype, 24% for family risk factor, and 5 years for age. Age prediction in groups including MCI and AD subjects involved several regions that were not prominent for age prediction otherwise. These regions included the middle and transverse temporal, paracentral and superior banks of temporal sulcus, as well as the amygdala and parahippocampal gyrus. The joint estimation of AD risk and connectome based mappings involved the cuneus, temporal, and cingulate cortices known to be associated with AD, and add new candidates, such as the cerebellum, whose role in AD is to be understood. Our predictive modeling approaches for AD risk factors represent a stepping stone towards single subject prediction, based on distances from normative graphs.

## 1 Introduction

While Alzheimer’s disease (AD) is thought of as a problem of old age, by the time it manifests clinically, it is too late for effective interventions for these individuals. Thus, we currently have no cure or means to arrest or reverse the course of AD. Identifying early biomarkers could open new avenues for implementing preventive lifestyle changes, or more effective interventions. One strategy to identify early biomarkers and targets for interventions is to study populations at risk, before cognitive decline. We have devised our study to include a sample from such populations, based on the major known genetic risk factor for late onset AD, the carriage of APOE4 alleles.

Structural connectomes hold untapped potential of revealing network based biomarkers, sensitive to cumulative effects from distinct pathologies, otherwise hard to detect. These vulnerable brain networks may be innate and genotype dependent, or reflect brain pathological changes, thought to exist long time before overt clinical AD manifestation and marked cognitive decline. While the problem of relating MR images or connectomes to traits is not new, it remains complicated because it involves large numbers of unknown parameters, relative to small study sample sizes. For example, the number of elements in an MRI image can reach 16777216 in a 256×256×256 linear voxel dimension cube, or in the case of connectomes it can be 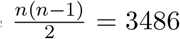 components of the lower triangular adjacency matrix for the widely used Desikhan-Killiany human brain atlas with n=84 regions.

Massive univariate analyses based on image/connectome components and a single trait may overfit the data, and suffer from lack of generalization ability. Moreover, univariate analyses relating imaging data to single predictors may not be adequate for the case of multifactorial diseases such as AD, and require considerations on predictor selection, and assessing the relationships between predictors [1]. Independent component analyses (ICA) and principal component analyses (PCA) approaches ([2, 3, 4, 5]) provide efficient ways to describe data through dimensionality reduction, and separating the signal from noise before regression analyses. However, these are unsupervised dimension reduction approaches. In addition, PCA projects the imaging data or connectomes to a subspace where all or some regions participate in a particular direction of interest, with different weights associated with them, hence they may lose the specificity of classical region of interest approaches. In contrast, sparse methods achieve higher specificity relative to PCA–like methods [6, 7], as well as taking into account another set of variable when reducing the dimension, i.e. supervised dimension reduction.

The Sparse Canonical Correlation Analysis (SCCA) method has been introduced and implemented in software packages (e.g. PMA, SCCAN). The results are mostly based on component estimation of coefficient weights. There has not been an attempt to extend the TNPCA methodology [8] with composite risk factors, in addition we use a bootstrap re-sampling method for constructing confidence intervals of the results of the SCCA. In this work we extended the TNPCA and the SCCA methodology to fill the mentioned gap, and used it in an application to brain connectome analysis.

Finally, we propose a predictive modeling approach that can move us closer to individualized subject prediction of AD risk, in asymptomatic individuals. Our predictive modeling framework is focused on age, and APOE genotype, as the primary risk factors for late onset AD. Our methodology can be easily expanded to other traits. While establishing relationships between brain structure and behavior has been a long–time goal in neuroscience, the majority of studies use functional or structural imaging, and more recently functional [9, 10] and structural connectomes [8]. Most studies do not perform a test on the accuracy or error of the model in a k-fold cross– validation scheme, which makes it difficult to asses the validity of such models for new data sets. Using connectome-based predictive modeling to predict individual behavior from brain connectivity is thus an important step in moving beyond simple correlation studies. Testing the performance in unseen samples quantifies the practical applicability of the derived models, while building accurate models remains a challenge [11].

Our methods provide stepping stones towards early identification of vulnerable brain networks, in relation to human traits relevant to AD risk. We illustrate the proposed framework in a set of subjects across the age span, and with various risk levels for AD, from control subjects with no known family history of AD, to subjects with known cases of AD in the family, to MCI and AD subjects.

## 2 Methods

In this work we developed three main threads: sparse canonical correlation with bootstrap confidence interval estimation (SCCA), Tensor Network SCCA (TNSCCA), and predictive modeling. We applied the proposed methods to a cohort of subjects with normal cognition, MCI and AD – to identify vulnerable brain networks associated with risk for AD, which can be expanded with a vector of human traits.

### 2.1 Sparse Canonical Correlation with Bootstrap Confidence Interval Estimation

Canonical correlation analysis (CCA) provides an avenue to jointly investigate relationships among multiple sets of data with different types. This attribute makes it particularly suitable for multifactorial diseases such as Alzheimer’s disease, where multiple pathologies e.g. amyloid, tau, neurodegeneration, may co-exist and interact to affect health and cognition of a subject, and thus may require the acquisition of different data types. CCA, and its different forms (constrained, multi set, nonlinear, sparse, etc) can be used to identify the common roots of statistical co–variations in multiple modalities, without assuming directionality. The reader is referred to [12] for an excellent review of studies using CCA in neuroscience. Here, we use CCA on structural connectomes derived from high resolution diffusion weighted magnetic resonance imaging, acquired at 3T, and on a constructed vector of AD risk factors, which we expanded with cognitive traits. The nature of this problem lends itself to a SCCA approach that can be used on images, as well as on connectomes.

Relative to CCA, SCCA considers the relationships between two types of variables but provides sparse solutions (coefficients), i.e. including only small subsets of variables of each type. SCCA maximizes the correlation between the projected subsets of variables via the estimated coefficients, while performing variable selection. The goal of SCCA is to identify linear combinations of the two sets of variables that are correlated with each other and associated with an outcome [13]. The first level SCCA method computes sparse projections vectors *u* ∈ ℝ^*p*^, *v* ∈ ℝ^*q*^ associated with *n* realization of multivariate random variables, 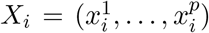 and 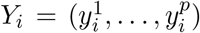 for *i* = 1, …, *n*, maximizing the correlation between the projected values 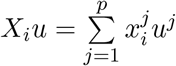 and 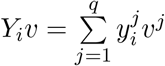.

To compute such projections, the sample level optimization problem is:

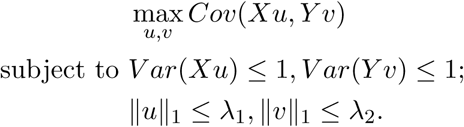

This problem has been studied in [14], where the regularization parameters *λ*_1_ and *λ*_2_ were tuned through a cross-validation procedure. We propose bootstrap re-sampling to make inference about the final result of this optimization problem, i.e. making inference about the projected correlation. In addition, since our studies included regions not usually visualized by current software programs, we extended the R brainconn [15] package by embedding the Desikan atlas with 84 regions of the human brain (IIT), and the cerebellum. This structure that was not present in the previous versions, has recently emerged to have a role in learning. This package is available at https://github.com/Ali-Mahzarnia/brainconn2.

We applied the proposed methods to identify vulnerable brain networks associated with risk for AD, including age, sex, and genotype. Both quantities, i.e. networks and traits, are attributed weights. Similar to most sparse regression models, there is no closed form solution to compute the p-value of the correlation between the projected values of the SCCA when testing whether it is zero in the population nor is there a formula to construct a confidence interval for such a population parameter. Hence, we use a re-sampling method to make inference about the projected correlation parameter in the population. There are two well–known re-sampling methods; bootstrapping and permutation. We choose bootstrapping to re-sample and make such an inference because it is supported by theoretical results of bootstrapping methods where the confidence interval of a statistics under bootstrap re-sampling approaches the true confidence interval in probability by increasing the sample size, without any prior assumption on its distribution, a.k.a consistency. Using the methodology of [16], we construct confidence intervals for the SCCA projected values correlation by re-sampling the data with replacement for 1000 times. After re-sampling and computing the 1000 correlations, we report the 2.5th and 97.5th percentiles of these correlations, i.e. the boundaries of the 95% confidence interval for population projected values correlation based on bootstrap percentiles. Since we cannot assume normality or symmetry of the distribution of these projected correlations [16] we cannot use a margin of error (normally distributed) method to construct symmetric confidence intervals.

### 2.2 Tensor Network SCCA

The Tensor network Principal components analysis (TNPCA) and its application to connectomes were introduced in [8]. This method first extracts *K* principal components of the tensors (brain connectomes) and uses them in a trait classification problem by Linear Discriminant Analysis (LDA) for a discrete binary response value trait, or in a trait prediction by CCA for a continuous univariate response value trait. Finally, this method uses the weights computed in the CCA or LDA to estimate the network changes associated with the response value variability.

We extend these methods in two different directions. The motivation behind such an extension is to reduce the dimension in a supervised manner while capturing the correlation between the projected values which reduces the data complexity. It is similar to performing PCA before running a sparse regression in a lower dimensions statistical analysis. First, instead of a univariate risk trait, we extend the results by incorporating a composite trait that is a multivariate random variable including both categorical and continuous random variables. Second, we use SCCA to compute sparse projections for the set of extracted TNPCA factors maximizing the correlation between the projected PC factors and the composite risk traits’ projected values.

TNPCA is an unsupervised dimension reduction statistical learning method on a set of connectomes, i.e. adjacency matrices 𝒳. The procedure is as follows:

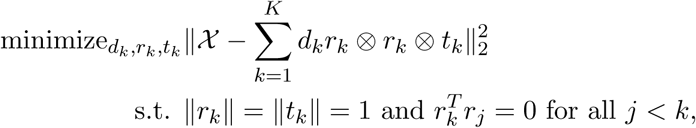

where *a* ⊗ *b* = *abT* is the outer product, *r*_*k*_ are vectors of p (number of nodes) and *t*_*k*_ are vector of *n* (sample size), that form the *k*_th_ CP decomposition factor and *d*_*k*_ is the kth positive CP scaling parameter. The CP method models a tensor X which is an array *p* by *p* by *n*, as a sum of rank one tensors. At this step, we deviate from the original TNPCA by using *K* factors *t*_*k*_ in a SCCA method coupled with a multivariate random variable *Y* as the composite trait.

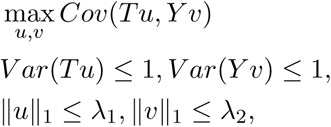

where *T* = (*t*_1_, …, *t*_*K*_) is a multivariate random variable of all *K* extracted factors by TNPCA. Next, similar to the original method, we measure the network change associated with the composite trait (or a linear combination of its components) by the formula that recovers the network based on *uk, tk, rk*, and *dk*:

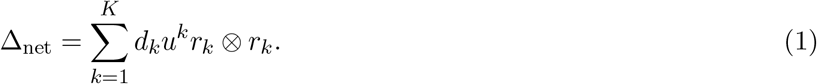

Then, for ease of interpretability we selected to report the top 30 connections that contributed to the measured network change, and thus are relevant to the composite risk factor variability in the sample.

One motivation behind TNSCCA method is the fact that the risk factor *Y* can be composite instead of only a continuous or a categorical (binary) variable, as in TNPCA. In addition, the principals are chosen based on their association with the risk factors in a supervised manner, while they are merely chosen based on their explanatory power of the tensors in TNPCA as an unsupervised dimension reduction method. Another important motivation is to reduce the dimensions for the networks associated to the risk traits via SCCA. Despite the fact that our study, having only 72 subjects, does not seem to be too large, such a method can help researchers with larger data sets and large sample sizes to reasonably reduce the dimension of the data while running SCCA efficiently.

### 2.3 Predictive modeling of AD risk traits

SCCA is fundamentally the same modelling approach as sparse regressions, such as LASSO and Elastic Net, given that one of the two variables is univariate. In this section we propose a predictive model for the SCCA when Y=(APOE genotype), Y=(family risk factor), and when Y=(Age) separately. We use the glmnet package for computation [17] with the following objective function,

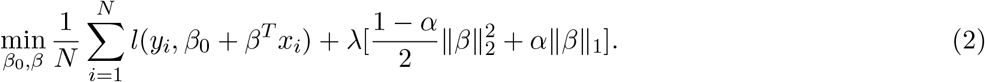

The third regression can be a sparse regression with continuous response value, i.e. 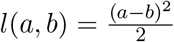, while the second is a sparse likelihood regression a.k.a sparse multinomial regression 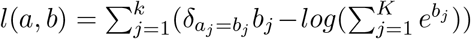 for *K* = 3 levels or groups and the vector Euclidean ∥∥ norms are replaced by the matrix Frobenius norm. In order to show the efficiency of such a method we compute the root mean squared error (RMSE) and average accuracy throughout k-fold (k=10) segmentation of data to train and validation sets respectively for each of the two regressions. We add an 0 *< α <* 1 grid search of 100 equally spaced values to the *λ*–tuning procedure of the glmnet package during the k–fold cross–validation. Finally, we run these regressions on the whole set to find the associated connectomes to APOE genotype, family risk, and age variability within the sample. Due to the small sample size, and the fact that there are not multiple models to be compared, it suffices to use this traditional k–fold analysis instead of designating a hold-out test set as often used for the final comparison between different models.

### 2.4 Experiments

#### 2.4.1 Subjects

All experimental procedures in this study were approved by the Duke University Medical Center Institutional Review Board. All participants, and where applicable their caretakers, provided informed consent, and were compensated for their time at the end of the study. 72 subjects were recruited, 35 males, and 37 females; age range 21-83; median age 50.5, average ± std: 51.22± 15.3. Age, sex, and genotype distribution for the study participants are shown in Table 1 and 2.

**Table 1:** Genotype and sex distribution for study participants

| Genotype / Sex | Male | Female |
| --- | --- | --- |
| APOE 23 | 3 | 3 |
| APOE 33 | 19 | 14 |
| APOE 34, 44 | 13 | 20 |

**Table 2:** Family risk and sex distribution for study participants. 0: subjects had no reported AD in the family, 1: subjects reported AD presence in family members; 2: mild cognitive impairment diagnosis; 3: AD diagnosis.

| Family risk / Sex | Male | Female |
| --- | --- | --- |
| 0 | 13 | 4 |
| 1 | 17 | 32 |
| 2: MCI (4), AD (2) | 5 | 1 |

We pooled MCI and AD subjects in the same AD risk category, as cognitively impaired subjects, due to small sample size of this subgroup in our subject populations.

#### 2.4.2 Image and Cognitive Data Collection

Cognitive testing was largely based on the Alzheimer’s Disease Centers’ Neuropsychological Test Battery in the Uniform Data Set (UDS3) [18]. The tests included: MOCA; Rey Auditory Verbal Learning Test (RAVLT) (immediate recall); Craft Story (immediate recall); Benson Figure Test (immediate score); Digit Span (forwards and backwards); Trail Making Test, Parts A and B; Digit Symbol; Categorical Verbal Fluency (animals, vegetables), Lexical Verbal Fluency (F-words, L-words); Benson figure test (delayed recall); Craft Story (delayed recall); RAVLT (delayed recall). The RAVLT Learning score was calculated as Trial 5 score minus the score of Trial 1, and RAVLT Forgetting as Trial 5 - Delayed Recall Score, for a list of 15 words. In addition we measured the useful field of view and the visual processing times needed to correctly identify objects in the presence of passive and active distractors (brainhq.com). Finally, we tested olfactory discrimination and olfactory memory. To measure olfactory memory we examined: a) encoding: 6 flavors were presented one-by-one, for 3 s each, separated by 30 s; b) recognition: a new set 6 odorants was presented after a 50 minutes delay, including new and previously presented stimuli. Participants were asked to rate, and label odorants. Odor ratings were measured on a 7-point Likert scale for: pleasantness: 1 (unpleasant)–7(pleasant); intensity: 1(weak)–7(strong); familiarity: 1(unfamiliar)–7(familiar); nameability: 1(not nameable) – 7(nameable). We measure the number and percentage of correctly recognized odorants scoring the number of correct answers from a binary choice whether the odorant presented in the second set was present or not in the first set (1(no)-7(yes)). We expect lower olfactory memory and discrimination with age and AD risk, with a dose dependent effect, increasing from normal to MCI and AD.

MRI was performed using a 3T GE Premier Performance scanner with Signa premier version 28, a 60 cm gradient coil with peak strength of 115 mT/m, and a 48-channel head coil. We acquired diffusion weighted images using motion robust, multi-shot echo planar imaging (EPI) with 4 interleaved shots, reconstructed using MUSE [19]. We used TE 60.6 ms/TR=14725 ms for 21 diffusion directions (b=1000 s/mm2) and 3 non-diffusion weighted images, acquired at 1×1×1 mm isotropic spatial resolution using a field of view (FOV) of 25.6 cm, and a 256×256 matrix, 136 slices, acquired in 24 minutes.

Anatomical images were used for brain parcellation and connectome estimation, and co-registration with DWI, and were acquired using a 3D T1-weighted IR-FSPGR-BRAVO sequence with TE=3.2 ms, TR=2263.7 ms, flip angle 8°, prep time 900 ms, recovery time 700 ms, 256×256×160 slices, 25.6×25×6×16 mm FOV, and image acquisition time was 5 min and 17 s.

#### 2.4.3 Preprocessing

For image analysis we used pipelines that we have developed for image registration, segmentation and voxel wise statistics, diffusion tensor estimation, tractography and connectomics [20, 21].

DWI processing used Dipy [22] to estimate the diffusion tensor and tractography. Reconstructed DWI volumes were affine registered to the first non-diffusion weighted image, using Advanced Normalization Tools (ANTs) [23], to correct for eddy distortions, and denoised using a PCA based algorithm [24].

All anatomical images were registered to the IIT human brain atlas [25] using the SAMBA [26] pipeline. A minimum deformation template was generated, to reduce individual subject biases. All subjects were mapped into the space of the minimum deformation template. The same registration was applied to the diffusion parameter maps previously obtained, and the reverse registration was used to bring the atlas labels into the subject space, providing 84 atlas labeled brain regions.

Structural connectomes were estimated as adjacency matrices, with each cell of the matrix containing the number of streamlines connecting a pair of atlas regions. As done in SCCA, prior to TNPCA we standardized the connectivity matrices component-wise across subjects. The streamlines were reconstructed using the Q-ball constant Solid Angle Reconstruction [27] implemented within Dipy, with the tractography results built using a deterministic maximum direction getter. We chose a relative peak threshold of 0.5, minimum separation angle at 25°, and a binary stopping criterion using the brain mask.

### 2.5 Modeling approaches

We used the methods introduced in 2.1, 2.2, 2.3 (bootstrap SCCA, TNSCCA, and predictive models respectively) on the data collected and pre–processed as described in section 2.4 to obtain the following results described in section 3.

Shared code: The codes are available at https://github.com/Ali-Mahzarnia/addecodemay2021 and the visualization R package is available at https://github.com/Ali-Mahzarnia/brainconn2. In addition, we have re–arranged the index of IIT atlas to better present its regions, in a machine readable format, appropriate for connectomics, and such index is available at https://github.com/Ali-Mahzarnia/atlasindex/blob/main/atlasindex.csv.

## 3 Results

We addressed three inter-related questions on identifying vulnerable brain networks associated with composite AD risk, and with performance in sensory and cognitive tests, as well as the ability to predict AD risk traits from connectomes. Traits were based on physiological and cognitive tests, i.e. the UDS3, including auditory, visual, and olfactory memory; and visual attention. For each analysis we have used as input the lower triangle of the connectivity matrices for each subject as *X*_*i*_ for *i* = 1, …, *n*, and a traits vector Y. We used the first step of SCCA for easier interpretations. Since SCCA is designed for the highest correlation between projected values, not the highest sparsity, we determined the regularization parameters *λ*_1_, *λ*_2_ via a net search for the risk traits, and the number of connectome projections respectively. SCCA and TNSCCA identified subnetworks associated with composite AD risk traits. Similarly we used sparse regression to predict categorical, e.g. APOE genotype and AD family risk, and continuous AD risk traits e.g. age, based on connectomes, and evaluated prediction performance.

### 3.1 Vulnerable brain networks associated with composite AD risk

#### 3.1.1 Composite AD risk: age, genotype, sex

Since old age, APOE4 genotypes and female sex are considered to increase risk for AD, we defined a composite AD risk factor as *Y* = (Sex, Genotype, Age). The risk component weights are reported in Table 3, and the associated brain subnetworks in Figure 1.

**Table 3:** AD risk traits and associated weights

| Factor | CCA weight |
| --- | --- |
| Sex | $-5.102 \times 10^{-3}$ |
| Age | $7.133 \times 10^{-2}$ |
| Genotype | $9.97 \times 10^{-1}$ |

**Figure 1:**
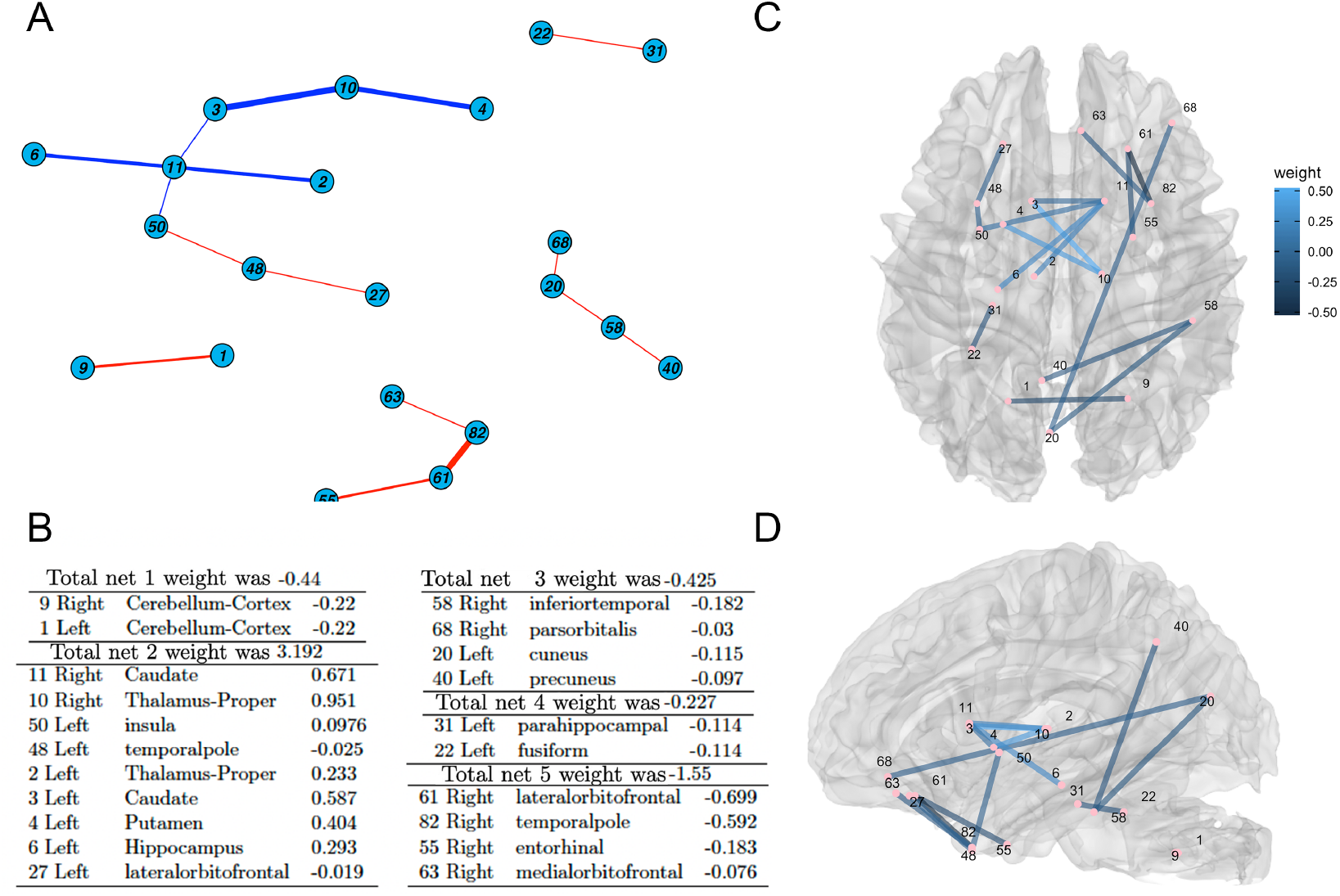
Vulnerable brain networks associated with the AD risk factor, defined as [age, genotype, sex]. A. 2D structure of the subgraphs, showing negative weights in red, to indicate pathology, and positive weights in blue. B. We identified 5 subnetworks, their anatomical region components, and estimated their weights. C. 3D rendering of brain subnetworks with reference to the brain anatomy (C: axial view, D: sagittal view)

There were 5 subnetworks associated with the AD risk factor, and 16 nonzero projected weights (Figure 1 A, B), from the 3486 components of the lower triangle of the connectome. The brain sub-graphs are shown in reference to the anatomy in Figure 1 C, D. The final correlation of the projected values was 0.735. The 95% confidence interval of the projected correlation based on 1000 bootstrap resamples was (0.653, 0.860).

Figure 2 shows that the resulting sparse networks separated groups by family risk, although this was not part of the variable in the analysis (Y). There was also separations by age; and by genotype, but not by sex. These plots serve only as illustration of detailed sub-network differences across subjects identified by the global SCCA. We also observed hyper–connectivity in subjects thought to represent higher risk, possibly due to over–compensation and different innate wiring; see e.g. sub-network 3 for the sex analysis.

**Figure 2:**
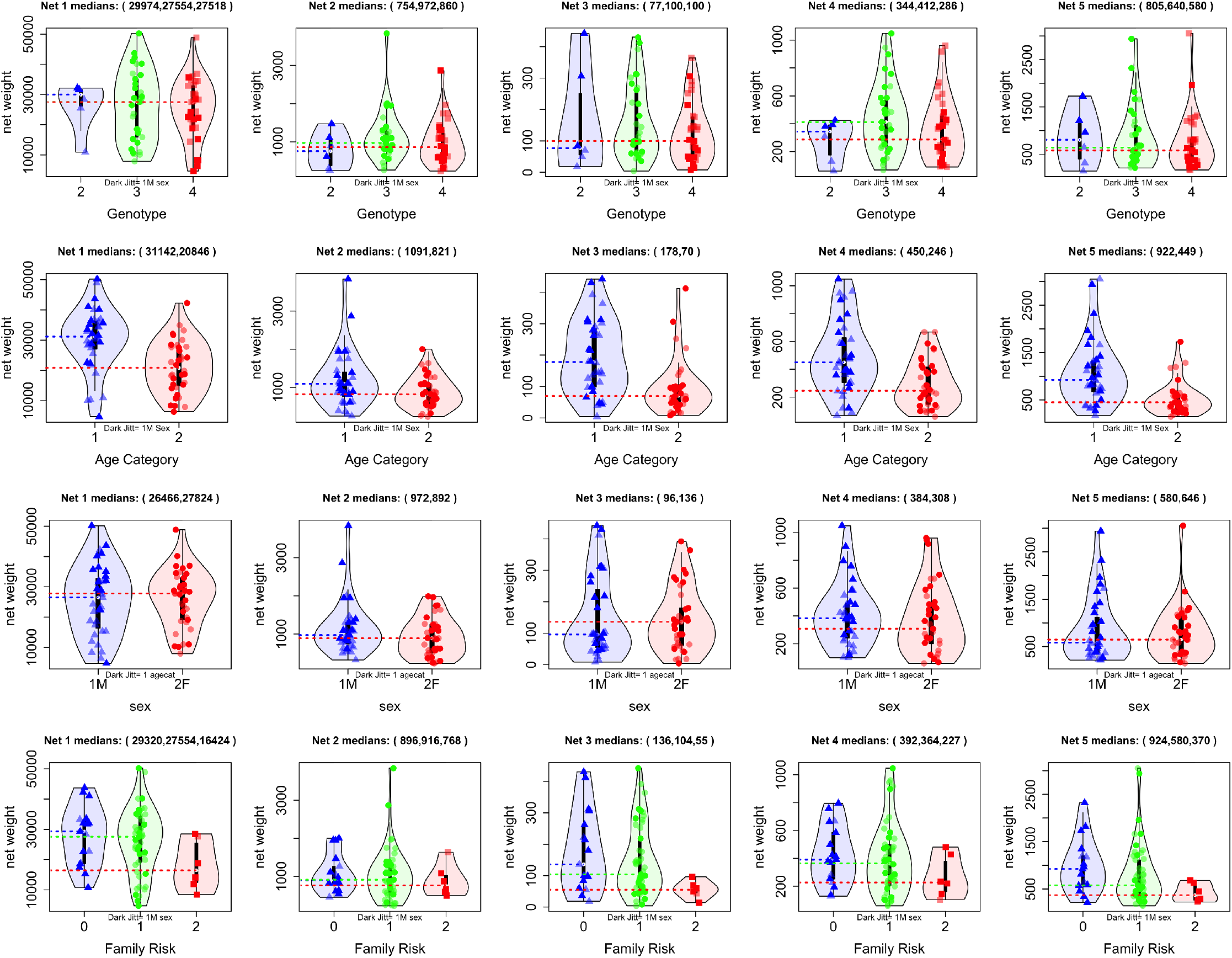
AD risk associated subnetworks connectivity by APOE genotype, age, sex, and AD family risk.

Our results revealed subgraphs associated with AD risk, and an important role for the APOE genotype (Table 3) in the composite AD risk. In general, higher AD risk corresponded to lower connectivity for sub-networks characterised mostly by negative edges (i.e. the opposite sign of age coefficient). The subgraph with the largest weight included the hippocampus, temporal pole and thalamus, as well as the insula, caudate, putamen and lateral orbito frontal cortex. Other subgraphs included the entorhinal cortex, precuneus and cuneus, as well as the cerebellum.

### 3.2 Mapping cognitive traits onto vulnerable networks

We estimated subnetworks associated with a subset of the UDS3 battery, including the Benson figure copy, RAVLT, and supplemented with olfactory memory, and visual attention testing, through the useful field of view (Figure 3).

**Figure 3:**
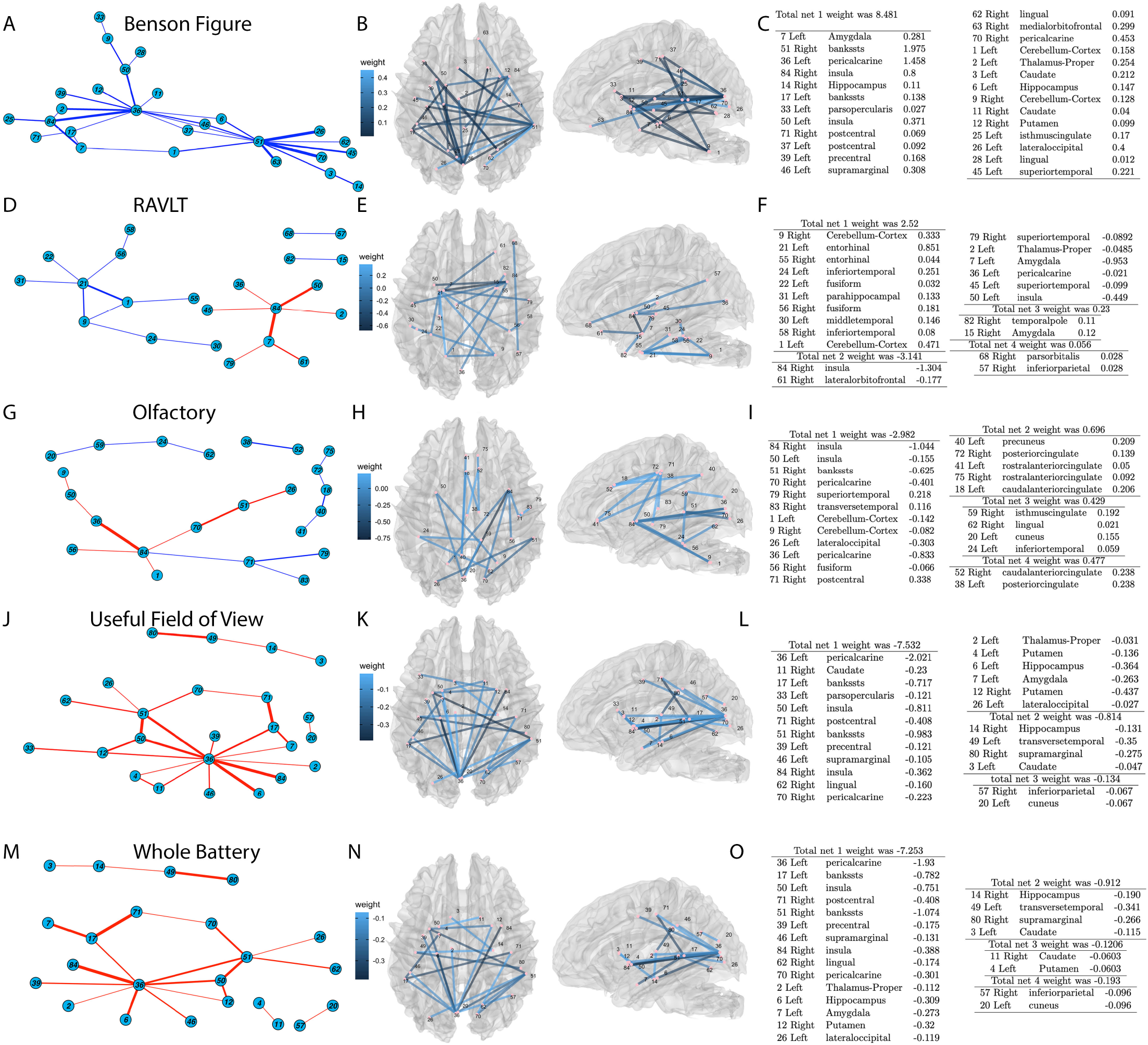
Brain networks associated with cognitive behavioral traits derived from: A-C. the Benson Figure Copy; D-F. RAVLT; supplemented with olfactory memory and; G-I. the useful field of view testing J-L; the whole testing battery; M-O.

#### 3.2.1 Benson Complex Figure Copy

The Benson figure test assesses visuo-constructional ability and visual memory, through copying (immediate memory) and recall (delayed memory) tests. We constructed a visual memory index Y=(Immediate Benson Score, Delayed Benson Score). The Y associated weights are reported in Table 4, and the X associated subnetworks in Figure 3 A-C. There were 31 nonzero projected weights associated with the connectomes. The final correlation of the projected values was 0.679, and its 95% bootstrap confidence interval was (0.64, 0.862).

**Table 4:**
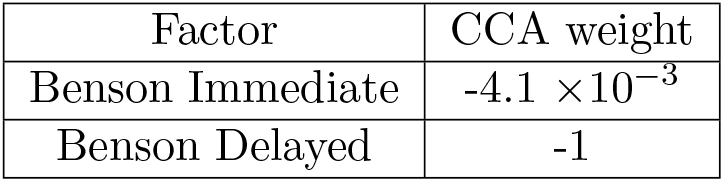
Benson figure drawing scores for the immediate and delayed components and their associated weights

The delayed Benson score had higher weight than the immediate score, which is practically 0. We identified one subnetwork, whose regions were involved in processing visual stimuli, e.g. the pericalcarine gyrus, containing the primary visual cortex, the insular cortex, also involved in visual perception, the lingual gyrus, and visuo-spatial orientation and attention (isthmus cingulate). This large subgraph also included areas involved in memory processing e.g. the hippocampus, amygdala, and medial prefrontal cortex, as well as somato-motor cortices, and multisensory integration cortices (superior temporal gyrus).

#### 3.2.2 Rey Auditory Verbal Learning Test - RAVLT

RAVLT tests auditory learning and memory, and we constructed a risk factor based on a subset of scores, *Y* = (RAVLT Immediate, RAVLT Learning, RAVLT Forgetting). The Y associated weights are reported in Table 5, and the subnetworks are shown in Figure 3 D-F. There were 19 nonzero projected weights associated with the 3486 components of the lower triangle of the connectome. The final correlation for the projected values was 0.652. The 95% bootstrap confidence interval of the projected correlation was (0.633, 0.841).

**Table 5:** RAVLT components associated weights

| Factor | CCA weight |
| --- | --- |
| RAVLT Immediate | 0.331 |
| RAVLT Learning | 0.944 |
| RAVLT Forgetting | $-7.248 \times 10^{-3}$ |

We identified four subgraphs associated with RAVLT performance. Three of these graphs had positive weights, while one had negative weights. This indicates possible compensatory mechanisms. The largest weight was associated with a connection for the insula, associated with auditory and visual perception, and the latero orbitofrontal cortex, involved in sensory integration and connected with limbic structures involved in emotion memory. The subgraph with the second largest weight included the middle temporal gyrus that subserves language and semantic memory processing, visual perception, and multimodal sensory integration, the entorhinal cortex involved in memory function; as well as the inferior temporal regions that respond to objects in complex scenes, and visual recognition in general; the fusiform gyrus involved in higher processing of visual information, including the identification and differentiation of objects, as well as in memory, multisensory integration and perception. Interestingly, the cerebellum was also part of this subgraph, supporting its role in sensory data acquisition and processing.

#### 3.2.3 Olfactory Memory

Since olfactory discrimination and memory may be early indicators of AD we constructed Y=(Composite Pleasantness, Composite Intensity, Composite Familiarity, Composite Nameability, Percent Correct Recall, Percentage Recognized). We noted the high correlations between these components (Table 6), which may be conducive to lower weights for correlated components (Table 7).

**Table 6:**
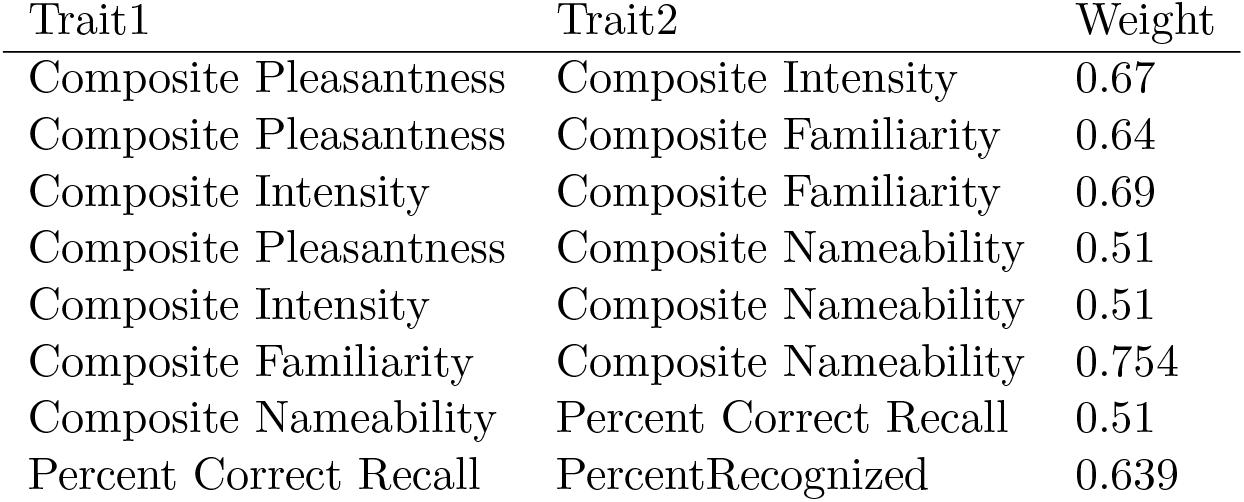
The olfactory memory predictors correlations

| Trait1 | Trait2 | Weight |
| --- | --- | --- |
| Composite Pleasantness | Composite Intensity | 0.67 |
| Composite Pleasantness | Composite Familiarity | 0.64 |
| Composite Intensity | Composite Familiarity | 0.69 |
| Composite Pleasantness | Composite Nameability | 0.51 |
| Composite Intensity | Composite Nameability | 0.51 |
| Composite Familiarity | Composite Nameability | 0.754 |
| Composite Nameability | Percent Correct Recall | 0.51 |
| Percent Correct Recall | PercentRecognized | 0.639 |

**Table 7:** The olfactory discrimination and memory components weights

| Factor | CCA |
| --- | --- |
| Composite Pleasantness | 0 |
| Composite Intensity | 0.247 |
| Composite Familiarity | 0.799 |
| Composite Nameability | 0.539 |
| Percent Correct Recall | 0.106 |
| Recognized | 0 |

There were 19 nonzero projected weights associated with 3486 components of the lower triangle of the connectomes matrix. The resulting networks are shown in Figure 3 G-I. The final correlation of the projected values was 0.704. The 95% bootstrap confidence interval of the projected correlation was (0.515, 0.844).

Among the measured scores, the odor familiarity and nameability had higher CCA values.The subgraph with the largest weight included the insula, involved in taste and smell processing; the post central gyrus - the primary somatosensory cortex; and the temporal and cingulate cortices.

#### 3.2.4 Useful Field of View - UFOV

UFOV measures visual processing speed and attention. We constructed Y based on Divided attention (UFOV2), and Selective attention (UFOV3) *Y* = (UFOV2, UFOV3), and found they had a high correlation of 0.8. The SCCA weights are reported in (Table 8), and there were 26 non-zero connectome components weights (Figure 3 J-L). The SCCA projected correlation was 0.715, and its 95% bootstrap confidence interval was (0.689, 0.894).

**Table 8:** Weights for UFOV trait components

| Factor | CCA |
| --- | --- |
| UFOV2 | -0.999 |
| UFOV3 | -0.004 |

The subgraphs with the highest weights included the percicalcarine areas, and insula with roles in vision and attention; the pars opercularis, shown to be thinner in subjects with ADHD [28]; the postcentral gyrus, is involved in proprieception and focal attention [29]; the Percentral gyrus, likely involved in the motor execution of the required task; the superior sulcus areas, that have been associated with joint attention (auditory/gaze) [30], and the supramarginal gyrus that has been associated with motor attention [31], and shifts of spatial attention [32].

#### 3.2.5 Mapping all traits together

We constructed a composite vector based on all measured traits, as Y=(Sex, APOE genotype, Systolic Pressure, Diastolic Pressure, Pulse, Height, Weight, BMI, Age, Family risk for AD, MOCA score, Benson Immediate, Benson Delayed, Percent Correct Recall, Percent Recognized, RAVLT Trial7, RAVLT Immediate, RAVLT Learning, RAVLT Forgetting, Fwd Digits total correct, Fwd Max Digit Length, Backwards Digits Total Correct, Backwards Digits Max Length, Trails A, Trails B, Trail Difference, Number Symbols, 4 Letter Fluency, Letter Fluency, UFOV2, UFOV3).

The correlations between the individual risk traits with absolute values more than 0.4 are reported in Table 9, and support that some of the measured traits may be dependent and thus redundant.

**Table 9:**
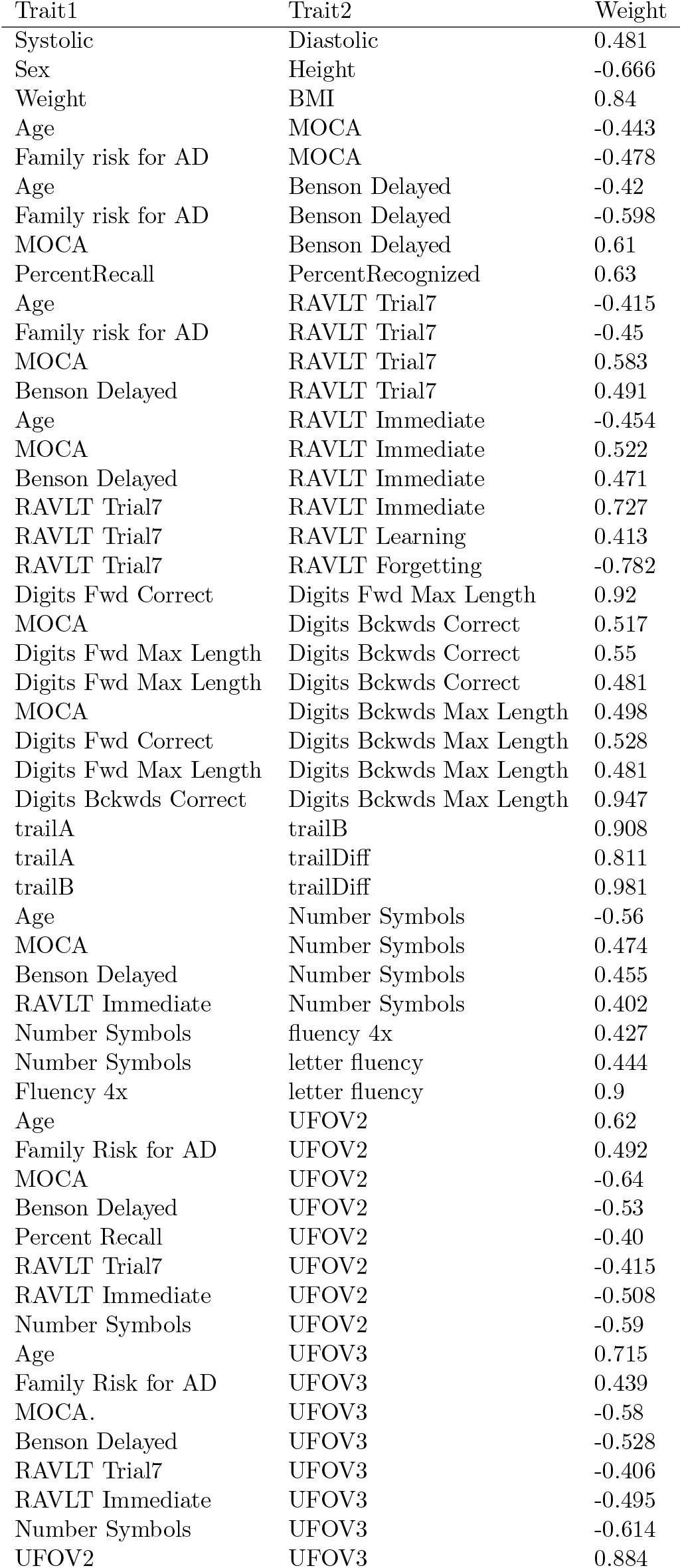
Correlation amongst traits included in the Y vector, selected if exceeding 0.4 in absolute value

The weights associated with the Y components are shown in Table 10. There were 22 nonzero projected weights associated with 3486 components of the lower triangle of a connectome matrix, and their weights are shown in Figure 3 O. The brain networks are shown in Figure 3 M-N. The final correlation of projected values was 0.735. The 95% bootstrap confidence interval of the projected correlation was (0.630, 0.894).

**Table 10:** The SCCA results for the combined measured traits (UDS3 and others) yielded a small set of variables with non-null weights

| Factor | CCA weight |
| --- | --- |
| Family Risk for AD | -0.18657 |
| MOCA | 0.564514 |
| Benson Delayed | 0.272723 |
| Number Symbols | 0.073921 |
| UFOV2 | -0.66472 |
| UFOV3 | -0.3533 |

Most networks have negative weights, therefore in those sub-nets a high connectivity can imply higher risk and a possible over-compensation or different innate wiring. This can happen e.g. for a study where the risk factor weight is also negative. For instance, a high a MOCA value indicates low risk. We note the large weight associated with the pericalcalrine, insula and temporal cortex structures (e.g. the banks of Superior Temporal Sulcus involved in social cognition, hippocampus, amygdala).

Integrating across these studies, we found the frequency of occurrence for a particular node was highest for the pericalcarine cortex, insula (3 times), and cerebellum; followed by the superior temporal gyrus, the banks of the superior sulcus, the fusiform gyrus, lingual cortex and amygdala.

Similarly, we examined the cumulative degree of connectivity of each region across the studies. Across the three cognitive studies, the regions with the highest cumulative degree of connectivity were the pericalcarine, insula, and the banks of the superior sulcus (*>* 10 edges), followed by the cerebellum, amygdala (6 edges), entorhinal cortex (5 edges), inferior temporal and postcentral gyri (4 edges). When adding the risk for AD the ranking remained similar, with the pericalcarine, insular and banks of superior sulcus at the top (*>* 13), followed by cerebellum and amygdala (7 edges), caudate, thalamus and entorhinal cortex (≥4), then hippocampus, cuneus, precuneus, fusiform gyrus, inferior temporal and latero orbital frontal gyri (3 edges).

### 3.3 TNSCCA Mapping vulnerable networks associated with AD risk

We extracted *K* = 80 principal components from the connectivity matrices during TNPCA; and all these factors together explained 96.8 of the variability in our data. We use these PCs as one random variable for SCCA, and we considered the other random variable to be Y=(sex, age, genotype, family risk factor). The associated weights are shown in Table 11 and Figure 4. The SCCA resulted in a projected correlation of 0.73. The 95% bootstrap confidence interval of the projected correlation based on 1000 re-samplings was (0.463, 0.810).

**Table 11:** Weights for the traits components used in TNSCCA

| Factors | CCA |
| --- | --- |
| sex | -0.008 |
| age | 0.751 |
| genotype | 0.285 |
| risk for ad | 0.594 |

**Figure 4:**
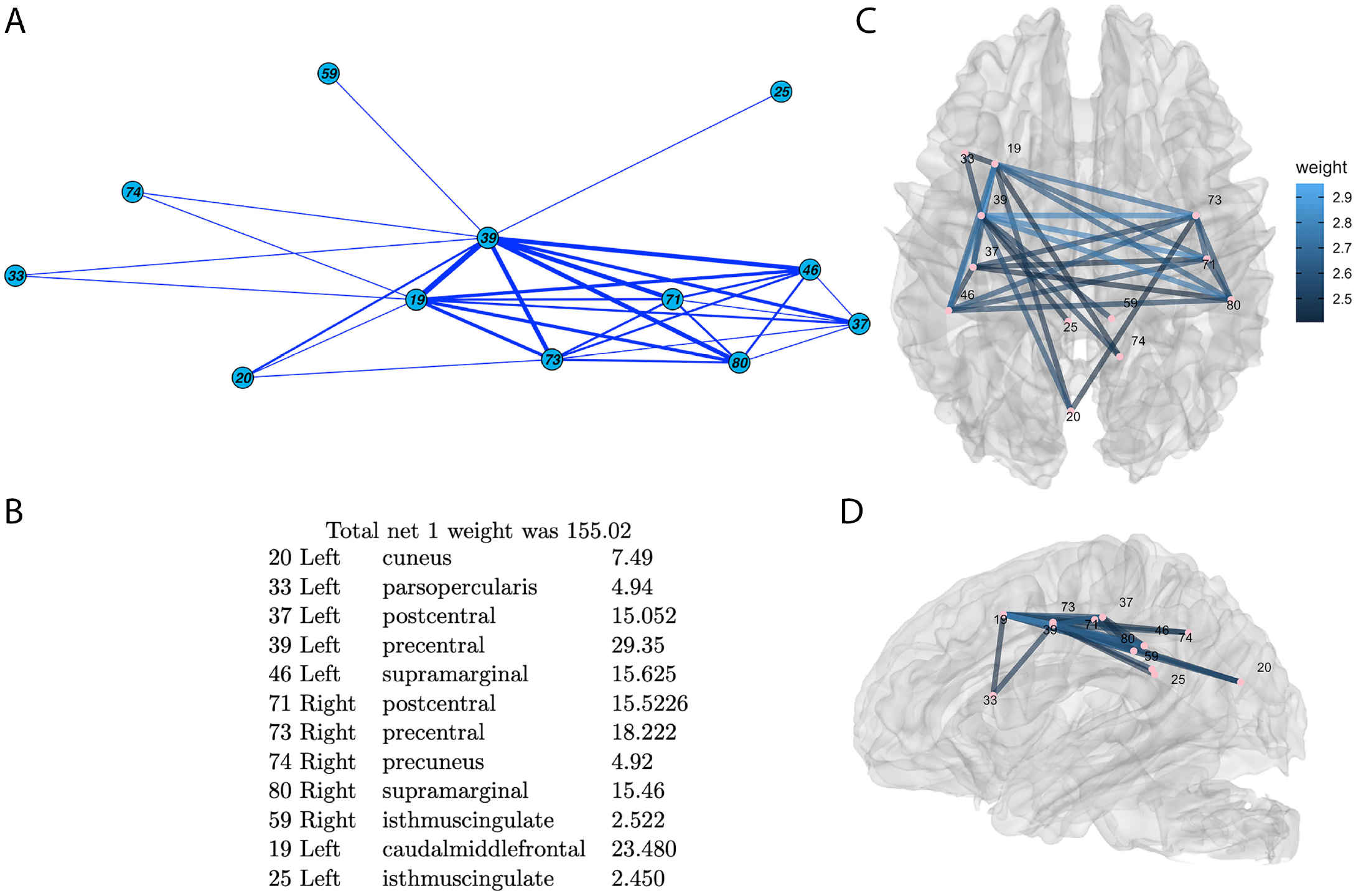
Vulnerable brain networks associated with AD risk, defined as [age, genotype, sex, family riskk (history of AD) ]. A. 2D structure of the 12 node subgraph, showing negative weights in red to indicate pathology, and positive weights in blue. B. The 5 subnetworks, their anatomical region components, and their weights. C-D. 3D rendering of brain subnetworks with reference to the brain anatomy (C: axial view, D sagittal view).

Age was the most significant component, followed by family risk, and genotype, while sex ranked last. The following principal factors were identified with non-zero coefficients in the SCCA.

We retrieved the network and measured the top 30 connections associated with the net variability, the Δ_net_ componenents are shown in Table 13.

**Table 12:** TNSCCA principal components and CCA weights

| PC | CCA |
| --- | --- |
| 2 | -0.82315 |
| 4 | -0.06774 |
| 5 | 0.355218 |
| 9 | -0.0247 |
| 13 | 0.222791 |
| 16 | -0.1869 |
| 23 | 0.317654 |
| 26 | -0.03691 |
| 58 | 0.064939 |

**Table 13:** Top connections and weights identified by TNSCCA when discriminating with a composite AD risk factor that includes age, sex, genotype and family history.

| Component1 | Hemisphere | Component2 | Hemisphere | Weight |
| --- | --- | --- | --- | --- |
| 'Percentral' | 'Left' | 'caudalmiddlefrontal' | 'Left' | '2.9572' |
| 'Percentral' | 'Right' | 'Percentral' | 'Left' | '2.8171' |
| 'supramarginal' | 'Left' | 'Percentral' | 'Left' | '2.8111' |
| 'postcentral' | 'Right' | 'Percentral' | 'Left' | '2.7992' |
| 'supramarginal' | 'Right' | 'Percentral' | 'Left' | '2.762' |
| 'Percentral' | 'Right' | 'caudalmiddlefrontal' | 'Left' | '2.7016' |
| 'supramarginal' | 'Left' | 'caudalmiddlefrontal' | 'Left' | '2.662' |
| 'Percentral' | 'Left' | 'postcentral' | 'Left' | '2.6537' |
| 'supramarginal' | 'Right' | 'caudalmiddlefrontal' | 'Left' | '2.635' |
| 'postcentral' | 'Right' | 'caudalmiddlefrontal' | 'Left' | '2.6128' |
| 'Percentral' | 'Right' | 'supramarginal' | 'Left' | '2.6057' |
| 'Percentral' | 'Right' | 'postcentral' | 'Right' | '2.5904' |
| 'Percentral' | 'Left' | 'cuneus' | 'Left' | '2.5813' |
| 'supramarginal' | 'Right' | 'Percentral' | 'Right' | '2.5767' |
| 'postcentral' | 'Left' | 'caudalmiddlefrontal' | 'Left' | '2.5693' |
| 'postcentral' | 'Right' | 'supramarginal' | 'Left' | '2.5549' |
| 'supramarginal' | 'Right' | 'supramarginal' | 'Left' | '2.5315' |
| 'supramarginal' | 'Right' | 'postcentral' | 'Right' | '2.5286' |
| 'isthmuscingulate' | 'Right' | 'Percentral' | 'Left' | '2.5227' |
| 'Percentral' | 'Right' | 'postcentral' | 'Left' | '2.5108' |
| 'precuneus' | 'Right' | 'Percentral' | 'Left' | '2.4987' |
| 'Percentral' | 'Left' | 'parsopercularis' | 'Left' | '2.4956' |
| 'cuneus' | 'Left' | 'caudalmiddlefrontal' | 'Left' | '2.4824' |
| 'supramarginal' | 'Left' | 'postcentral' | 'Left' | '2.4597' |
| 'Percentral' | 'Left' | 'isthmuscingulate' | 'Left' | '2.4508' |
| 'parsopercularis' | 'Left' | 'caudalmiddlefrontal' | 'Left' | '2.4416' |
| 'postcentral' | 'Right' | 'postcentral' | 'Left' | '2.4368' |
| 'supramarginal' | 'Right' | 'postcentral' | 'Left' | '2.4224' |
| 'Percentral' | 'Right' | 'cuneus' | 'Left' | '2.4201' |
| 'precuneus' | 'Right' | 'caudalmiddlefrontal' | 'Left' | '2.4189' |
| 'isthmuscingulate' | 'Right' | 'caudalmiddlefrontal' | 'Left' | '2.4178' |

TNSCCA identified the top connections associated with a composite AD risk factor that included age, sex, genotype and family history. This rather large network contained the precuneus-precentral gyrus connections, and precentral and postcentral interhemishperic connections. Notably the precuneus has been shown to be involved in recollection and memory, integration of information relating to perception of the environment, cue reactivity, and episodic memory retrieval.

### 3.4 Predictive models

#### 3.4.1 APOE Genotype

We used elastic net for predicting APOE genotype, and a 10 fold cross validation strategy for tuning the two parameters *α* and *λ* in a net. The miss-classification rate for predicting APOE genotype over the whole set was 16.7%. The final result had only 9 non-zero coefficients associated with the lower triangle of the connectomes (Figure 5).

**Figure 5:**
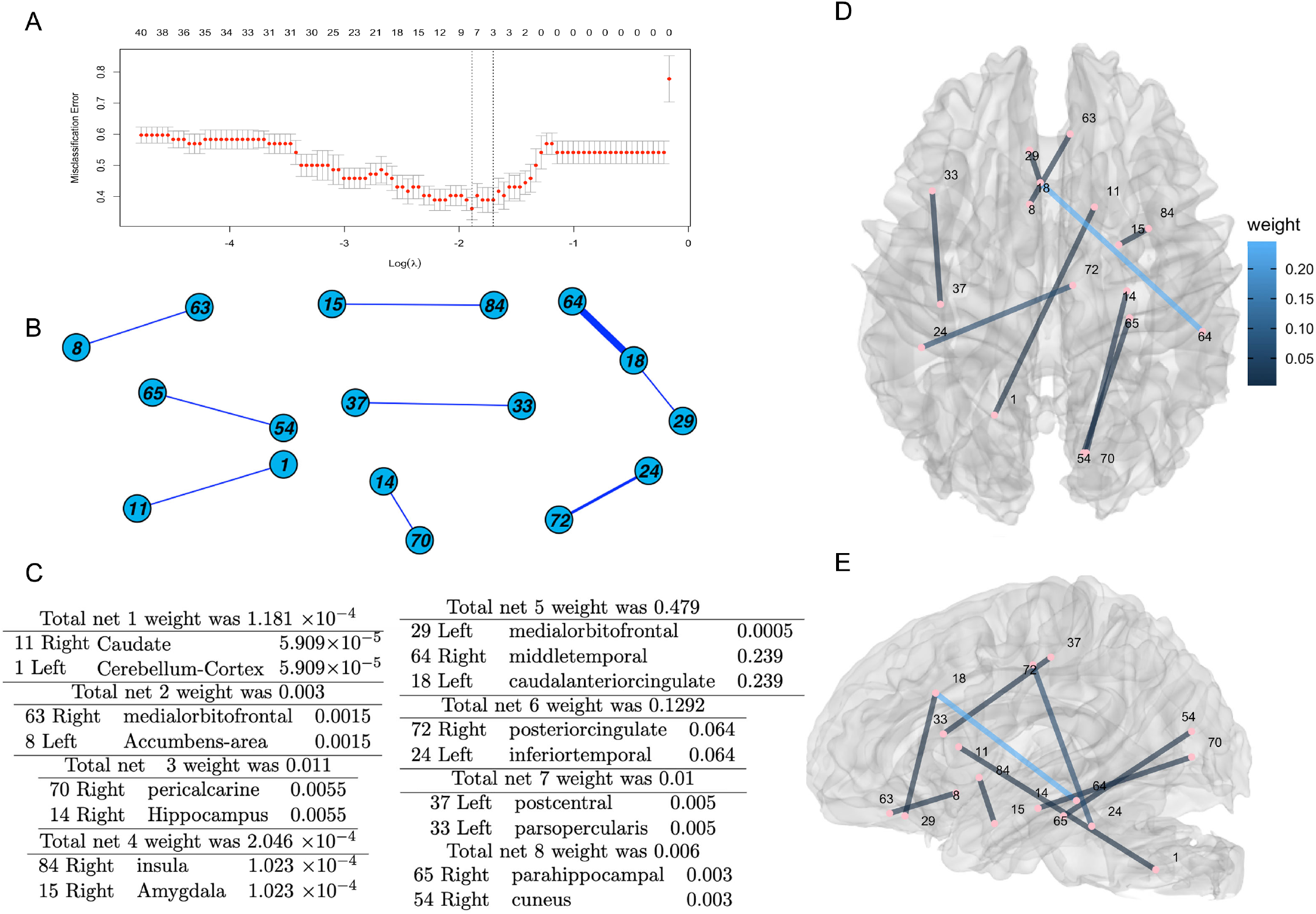
Brain subnetworks predictive of genotype. A. Parameter selection was based on minimizing the missclassification error. B 2D structure of the subgraphs. C. We identified 7 subnetworks, their anatomical region components, and associated weights. D-E. 3D rendering of brain subnetworks with reference to the brain anatomy (D: axial view, E sagittal view).

We identified brain connections predictive of APOE genotype, including those between the posterior cingulate and the inferior temporal lobe, as well as other regions involved in memory, the cuneus and parahippocampal gyrus, but also the caudate, and cerebellum.

#### 3.4.2 Family risk factor

We used Elastic Net for predicting the family risk factor, and a *k* = 10–fold cross-validation strategy for tuning the two parameters (Figure 6). The miss-classification rate over the whole set was 23.6%. We identified 9 non-zero coefficients associated with the components of the lower connectivity matrices. A sparse subgraph was associated with the family risk factor, and the nodes with the largest total weight included the traverse temporal, accumbens, frontal pole, and caudal anterior cingulate.

**Figure 6:**
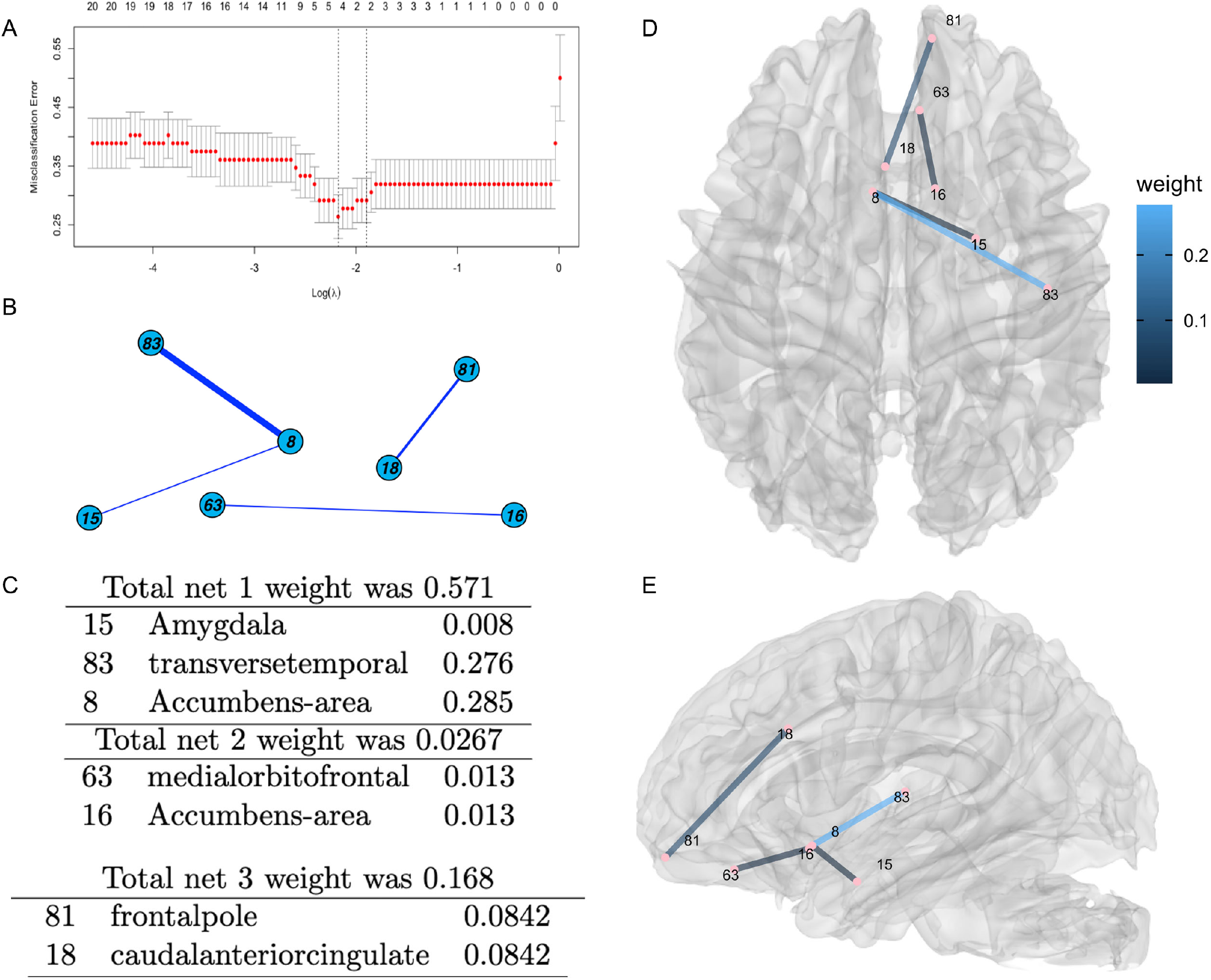
Brain networks predictive of family risk for AD. A. Parameter selection based on minimizing missclassification error based on kfold average results. B 2D structure of the identified subgraphs. C. The 3 subnetworks, their anatomical region components, and associated weights. D-E. 3D rendering of brain subnetworks with reference to the brain anatomy (D: axial view, E sagittal view).

#### 3.4.3 Age

When using the Elastic Net, the RMSE was 5.324 years for predicting age for the whole set after tuning the two parameters through 10–fold cross-validation. The standard deviation for age was 15.301 years (Figure 7). Hence, a considerable amount of sample variability (67%) of age can be explained by such a sparse linear relationship. The final result has 49 non-zero coefficients associated with the components of the lower connectivity matrices.

**Figure 7:**
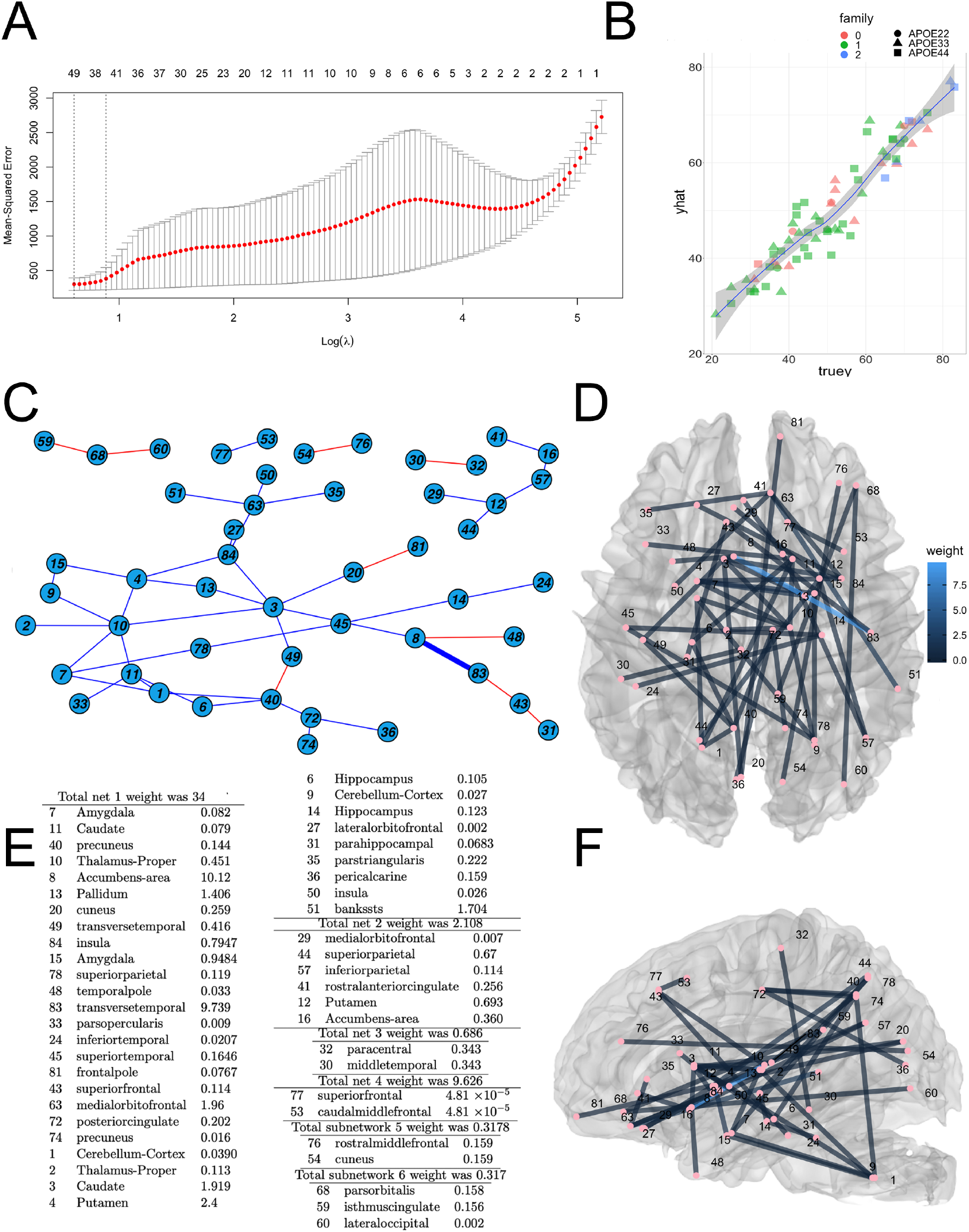
Age prediction based on connectomes; A. Age missclassification error based on kfold average results; B. True age versus predicted age; C. 2D structure of the identified subgraphs. E. The 6 subnetworks, their anatomical region components, and associated weights. D and F. 3D rendering of brain subnetworks with reference to the brain anatomy (D: axial view, F: sagittal view).

As expected age prediction involved extensive brain networks, and the large subgraph was associated with the highest weight. A large weight was also found for the accumbens. Other large weight nodes in this subgraph included the transverse temporal gyrus, the banks of the superior temporal sulcus, the putamen, hippocampus and amygdala.

When predicting age without the MCI and AD groups we identified a smaller set of regions that when including this subpopulation. 12 connections were only present in the group with MCI/AD but had no role in normal age prediction. These regions included the middletemporal, paracentral, transversetemporal cortices and banks if the superior sulcus.

In summary we have developed approaches to associate vulnerable brain networks with traits, and predictive modeling approaches for genotype, family risk factor, and age. Our predictive modeling approaches will allow to compare populations at risk, across the age span, based e.g. on overestimation or underestimation of biological age, and have potential to reveal new targets based on vulnerable brain networks.

## 4 Discussion

One of the key questions in brain research seeks to identify brain networks that are vulnerable, and those that are resilient to aging and AD. Identifying network based biomarkers in populations at risk would allow for developing more effective strategies than currently available, either through preventive or therapeutic interventions.

We have focused our efforts on jointly identifying structural brain networks and associated risk factors for AD in people with APOE2, APOE3, and APOE4 genotypes. These three major APOE alleles are associated with protection for AD, control populations, and with increased genetic risk, respectively. We have used SCCA to identify vulnerable subgraphs and AD risk factors, and their relative contributions, through their CCA weights. Moreover we extended the CCA result for the projected correlation by associating it with bootstrap confidence intervals. These strategies provided insight for region contributions, and helped weigh predictors extracted from the proposed AD risk factors, either physiological, or based on cognitive traits.

The examination of AD risk associated networks and weights revealed a role for a subnetowrk including the temporal pole, lateral orbital frontal cortex, and entorhinal cortex. A large literature exist regarding AD pathology in the entorhinal cortex [33]. The temporal pole is known to have a significant role in recognizing and using language, but it also helps with object recognition, and creating new and long term memories, and suffers changes in AD [34]. The lateral orbital frontal cortex has also been found to be damaged in AD, and may help explain the underlying mechanisms for behavioral changes accompanying AD, including sensory-related processes i.e. taste and olfaction, decision making, mood, social behavior and aggressiveness in personality [35]. The second subgraph in terms of total weight included the interhemispheric connectivity of the cerebellum. While its role in AD has not been fully explored, the cerebellum has string cerebral connections, e.g. to the prefrontal and parietal cortices, the cingulate and parahippocampal gyri [36, 37, 38]. These connections support a substrate for cerebellar involvement in cognitive processes, which is yet to be fully understood. The cerebellum receives input from the inferior olivary nuclei and the locus coeruleus, contributing to sensorimotor and memory functions [39, 40]. We note that the locus coeruleus is thought to be the initial site of tau pathology [41, 42, 43], which may potentiate amyloid pathology that only affects the cerebellum in later stages [44].

We have identified a role for cerebellar connectivity, suggesting that more investigations are needed to understand its modulatory role of the brain–behaviour relationship in neurodegenerative disorders. We note that many previous AD studies have excluded the cerebellum, and many software packages remove the cerebellum from consideration, which prompted us to modify existing packages to visualize the cerebellar-cerebral connectivity. Perhaps one of the challenges of including the cerebellum in previous studies stems from difficulties imaging these areas. Improved MRI hardware, with increased sensitivity and better shimming leading to increase signal and reduce magnetic field inhomogeneities, fuels the potential for future studies on this region, increasing the attention it deserves in aging studies.

The third network involved the inferior temporal gyrus, with a known role in visual object recognition and the cuneus, also involved in visual information processing. These findings support fMRI results on network connectopathies in AD, with a role for visual processing networks [45], and point toward the need to further explore sensory processing in aging and AD.

We have next mapped vulnerable circuits associated with visual, auditory, and olfactory memory, as well as with a subset of the UDS3 cognitive battery, complemented with physiological measures (e.g. on blood pressure, BMI). Integrating across these separate studies the frequency of occurrence was highest for the pericalcarine cortex, insula, and cerebellum (present in all three results for visual, auditory, and olfactory memory); followed by the superior temporal gyrus, the banks of the superior sulcus, the fusiform gyrus, lingual cortex and amygdala (present in two of the results). The hippocampus, entorhinal and cingulate cortices also appeared as regions, in at least one instance. Pericalcarine cortical thinning has been reported in aging and also AD [46], and interestingly in first degree relatives of AD patients [47], where the effect was largely attributed to APOE4 effects. The pericalcarine cortical thinning can also be linked to gait impairment [48]. The role of the insula can be linked to several functions, notably attention [49, 50] ; and also to olfactory. Olfactory function decline has been reported prior to cognitive and memory decline in AD. Our results support the role of olfaction as a source for potential early biomarkers [51]. The role of cerebellum in cognition and behavior has recently received increasing recognition [52]. The highest cumulative degree of connectivity (sum of total number of edges for a region) across all three cognitive studies plus the AD composite risk factor revealed a role for the pericalcarine, insular, and banks of superior sulcus with (*>*13 edges), followed by cerebellum (7 edges), amygdala (6 edges), caudate, thalamus and entorhinal cortex (4 edges), and then by the hippocampus, cuneus, precuneus, fusiform gyrus, inferior temporal and latero orbital frontal gyri (3 edges). These findings confirmed expectations of altered connectivity in visual, language, and olfactory areas respectively; and also indicated the presence of more extensive networks. These results support the role of white matter connecting remote gray matter areas responsible for cognitive processes, and just as above, that more investigations into sensory function alterations in AD are needed.

This study complements the larger literature that has sought to identify vulnerable based on functional MRI [53]. Our results linking structural connectome with AD risk and cognitive traits largely support and complement published fMRI network biomarkers studies, but in preclinical populations. Our findings underline giving more consideration to several potential early biomarkers, including a role for the cerebellum.

We have extended tensor network principal component analyses to include both categorical and continuous random variables, in this case reducing dimensions while capturing correlation. We used SCCA to select sparse projections for the set of extracted TNPCA factors (maximizing these projection correlation with the risk traits’ projected values). We have included age, genotype, sex, and family risk for AD on the Y side, and our results indicated a major role for age, followed by family risk, and genotype, while sex had a much smaller weight, compared to age. The top connections emphasised a role of the precentral gyrus, involved in motor control, and associated with normal aging processes [54]. Other prominent regions included the caudal middle frontal (involved in default mode network connectopathies in aging and cognitive disorders [55], postcentral [56], and supramarginal gyrus (showing cytoarchitectural changes in LOAD) [57], and the precuneus [58].

We identified brain subnetworks predictive of APOE genotype (16% error), AD family risk (24%), and age (5 years). Predicting a greater brain age than the biological one may help identify subjects that are cognitively normal but at risk for AD, and early targets that show changes before cognitive decline. Making such predictions in cognitively normal versus those with cognitive decline identified large sub networks, with overlap between the two groups, but also a small set of 12 connections that were revealed to be present in the group with MCI/AD but had no role in normal age prediction. These regions included the middletemporal, paracentral, transversetemporal cortices and banks if the superior sulcus.

Limitations of our study come from a small population sample, in particular for the cognitively impaired subgroup. We thus aim to replicate this study in a larger population, and possibly using existing data bases, which may inform on the validity of current findings. One significant challenge in this approach lies into harmonizing network data sets based on images acquired at different sites, with different diffusion schemes and spatial resolutions. Short of replication, new methods are needed to translate from one study to another. The other limitation is that while networks may be sensitive to the cumulative effects of pathologies preceding clinical AD, more studies using different imaging methods are needed to disentangle the precise mechanisms underlying the observed changes. Optimal predictor and model selection are other factors to be taken into consideration, and to refine future models.

Previous connectome based predictive modeling efforts have largely focused on one single trait prediction at a time [11]. In addition, the significant edges were selected based on numerous individual simple regression models. Our predictive model advances the field in that it takes into account the covariance structure within the connectome, in one sparse regression model. [11] used a hold-out test and obtained a correlation of 0.87 between the predicted and true trait values. Wile we did not compare different models, and did not have a hold–out set because of the small sample size, we obtained a correlation of 0.947 for age prediction.

Previous studies relying mostly on fMRI functional connectomes and Pearson correlation have identified brain regions and networks discriminating APOE4 carriers from controls, but there are relatively fewer studies focusing on preclinical stages [59], and on structural connectomes [60], which may confer increased robustness. Here we have used high resolution diffusion based imaging, but joint analyses [61, 60] of both structural and functional connectomes may provide advantageous, in particular for task specific network mapping. Such joint analyses may provide a better understanding of structure-function coupling [62], and the effects of disconnection or hyperconnection alterations as mechanistic substrates for selective circuit vulnerability or resilience in individuals with different APOE genotypes during aging and AD. These advances provide stepping stones towards genotype specific approaches addressing problems of aging and AD.

## 5 Acknowledgements

We are most grateful to the volunteers who have participated in this study, for their time, suggestions, and enthusiastic support. We are also thankful to the members of BIAC, the Duke Radiology, and Duke Neurology Departments for generously sharing advice, support and resources needed for the experiments, in particular to Iain Bruce and Todd Harshbarger for help setting up imaging protocols, to Chris Petty and Francis Favorini for help setting up, and maintaining the computational environment, to Susan Music, Lamont Conyers and Jennifer Graves for assistance with scanning. This work was supported by RF1 AG057895, R01 AG066184, U24 CA220245, RF1 AG070149, and P30 AG072958.

## References

[1] Priya Ranganathan, CS Pramesh, and Rakesh Aggarwal. Common pitfalls in statistical analysis: logistic regression. Perspectives in clinical research, 8(3):148, 2017.

[2] B Borroni, M Grassi, E Premi, S Gazzina, A Alberici, M Cosseddu, B Paghera, and A Padovani. Neuroanatomical correlates of behavioural phenotypes in behavioural variant of frontotemporal dementia. Behavioural brain research, 235(2):124–129, 2012.

[3] Pierre Comon. Independent component analysis, a new concept? Signal processing, 36(3):287–314, 1994.

[4] Peter Mansfield. Multi-planar image formation using nmr spin echoes. Journal of Physics C: Solid State Physics, 10(3):L55, 1977.

[5] Jul Lea Shamy, Christian Habeck, Patrick R Hof, David G Amaral, Sania G Fong, Michael H Buonocore, Yaakov Stern, Carol A Barnes, and Peter R Rapp. Volumetric correlates of spatiotemporal working and recognition memory impairment in aged rhesus monkeys. Cerebral cortex, 21(7):1559–1573, 2011.

[6] Daniel D Lee and H Sebastian Seung. Learning the parts of objects by non-negative matrix factorization. Nature, 401(6755):788–791, 1999.

[7] Brian B Avants, David J Libon, Katya Rascovsky, Ashley Boller, Corey T McMillan, Lauren Massimo, H Branch Coslett, Anjan Chatterjee, Rachel G Gross, and Murray Grossman. Sparse canonical correlation analysis relates network-level atrophy to multivariate cognitive measures in a neurodegenerative population. Neuroimage, 84:698–711, 2014.

[8] Zhengwu Zhang, Genevera I Allen, Hongtu Zhu, and David Dunson. Tensor network factorizations: Relationships between brain structural connectomes and traits. Neuroimage, 197:330–343, 2019.

[9] Ali Mahzarnia and Jun Song. Multivariate functional group sparse regression: Functional predictor selection. PloS one, 17(4):e0265940, 2022.

[10] Diana O Svaldi, Joaquín Goñi, Kausar Abbas, Enrico Amico, David G Clark, Charanya Muralidharan, Mario Dzemidzic, John D West, Shannon L Risacher, Andrew J Saykin, et al. Optimizing differential identifiability improves connectome predictive modeling of cognitive deficits from functional connectivity in alzheimer’s disease. Human brain mapping, 42(11):3500–3516, 2021.

[11] Xilin Shen, Emily S Finn, Dustin Scheinost, Monica D Rosenberg, Marvin M Chun, Xenophon Papademetris, and R Todd Constable. Using connectome-based predictive modeling to predict individual behavior from brain connectivity. nature protocols, 12(3):506–518, 2017.

[12] Xiaowei Zhuang, Zhengshi Yang, and Dietmar Cordes. A technical review of canonical correlation analysis for neuroscience applications. Human Brain Mapping, 41(13):3807–3833, 2020.

[13] Daniela M Witten and Robert J Tibshirani. Extensions of sparse canonical correlation analysis with applications to genomic data. Statistical applications in genetics and molecular biology, 8(1), 2009.

[14] Daniela M Witten, Robert Tibshirani, and Trevor Hastie. A penalized matrix decomposition, with applications to sparse principal components and canonical correlation analysis. Biostatistics, 10(3):515–534, 2009.

[15] Edwina R Orchard, Phillip GD Ward, Sidhant Chopra, Elsdon Storey, Gary F Egan, and Sharna D Jamadar. Neuroprotective effects of motherhood on brain function in late life: a resting-state fmri study. Cerebral Cortex, 31(2):1270–1283, 2021.

[16] Robert J Tibshirani and Bradley Efron. An introduction to the bootstrap. Monographs on statistics and applied probability, 57:1–436, 1993.

[17] Jerome Friedman, Trevor Hastie, and Rob Tibshirani. Regularization paths for generalized linear models via coordinate descent. Journal of statistical software, 33(1):1, 2010.

[18] Sandra Weintraub, Lilah Besser, Hiroko H Dodge, Merilee Teylan, Steven Ferris, Felicia C Goldstein, Bruno Giordani, Joel Kramer, David Loewenstein, Dan Marson, et al. Version 3 of the alzheimer disease centers’ neuropsychological test battery in the uniform data set (uds). Alzheimer disease and associated disorders, 32(1):10, 2018.

[19] Nan-kuei Chen, Arnaud Guidon, Hing-Chiu Chang, and Allen W Song. A robust multi-shot scan strategy for high-resolution diffusion weighted mri enabled by multiplexed sensitivity-encoding (muse). Neuroimage, 72:41–47, 2013.

[20] Robert J Anderson, James J Cook, Natalie Delpratt, John C Nouls, Bin Gu, James O McNamara, Brian B Avants, G Allan Johnson, and Alexandra Badea. Small animal multivariate brain analysis (samba)–a high throughput pipeline with a validation framework. Neuroinformatics, 17(3):451–472, 2019.

[21] Robert J Anderson, Christopher M Long, Evan D Calabrese, Scott H Robertson, G Allan Johnson, Gary P Cofer, Richard J O’Brien, and Alexandra Badea. Optimizing diffusion imaging protocols for structural connectomics in mouse models of neurological conditions. Frontiers in physics, 8:88, 2020.

[22] Eleftherios Garyfallidis, Matthew Brett, Bagrat Amirbekian, Ariel Rokem, Stefan Van Der Walt, Maxime Descoteaux, Ian Nimmo-Smith, and Dipy Contributors. Dipy, a library for the analysis of diffusion mri data. Frontiers in neuroinformatics, 8:8, 2014.

[23] Brian B Avants, Charles L Epstein, Murray Grossman, and James C Gee. Symmetric diffeomorphic image registration with cross-correlation: evaluating automated labeling of elderly and neurodegenerative brain. Medical image analysis, 12(1):26–41, 2008.

[24] Jelle Veraart, Dmitry S Novikov, Daan Christiaens, Benjamin Ades-Aron, Jan Sijbers, and Els Fieremans. Denoising of diffusion mri using random matrix theory. Neuroimage, 142:394–406, 2016.

[25] Shengwei Zhang and Konstantinos Arfanakis. Evaluation of standardized and study-specific diffusion tensor imaging templates of the adult human brain: Template characteristics, spatial normalization accuracy, and detection of small inter-group fa differences. Neuroimage, 172:40–50, 2018.

[26] Robert J Anderson, James J Cook, Natalie Delpratt, John C Nouls, Bin Gu, James O McNamara, Brian B Avants, G Allan Johnson, and Alexandra Badea. Small animal multivariate brain analysis (samba)–a high throughput pipeline with a validation framework. Neuroinformatics, 17(3):451–472, 2019.

[27] David S Tuch. Q-ball imaging. Magnetic Resonance in Medicine: An Official Journal of the International Society for Magnetic Resonance in Medicine, 52(6):1358–1372, 2004.

[28] Martin J Batty, Elizabeth B Liddle, Alain Pitiot, Roberto Toro, Madeleine J Groom, Gaia Scerif, Mario Liotti, Peter F Liddle, Tomáš Paus, and Chris Hollis. Cortical gray matter in attention-deficit/hyperactivity disorder: a structural magnetic resonance imaging study. Journal of the American Academy of Child & Adolescent Psychiatry, 49(3):229–238, 2010.

[29] Pa E Roland. Somatotopical tuning of postcentral gyrus during focal attention in man. a regional cerebral blood flow study. Journal of Neurophysiology, 46(4):744–754, 1981.

[30] Simone Materna, Peter W Dicke, and Peter Thier. Dissociable roles of the superior temporal sulcus and the intraparietal sulcus in joint attention: a functional magnetic resonance imaging study. Journal of cognitive neuroscience, 20(1):108–119, 2008.

[31] Matthew FS Rushworth, Philip D Nixon, Shelley Renowden, Derick T Wade, and Richard E Passingham. The left parietal cortex and motor attention. Neuropsychologia, 35(9):1261–1273, 1997.

[32] Timothy J. Silk, Mark A. Bellgrove, Pia Wrafter, Jason B. Mattingley, and Ross Cunnington. Spatial working memory and spatial attention rely on common neural processes in the intraparietal sulcus. NeuroImage, 53(2):718–724, 2010.

[33] Heiko Braak and Eva Braak. Neuropathological stageing of alzheimer-related changes. Acta neuropathologica, 82(4):239–259, 1991.

[34] GB Karas, Emma J Burton, Serge ARB Rombouts, Ronald A van Schijndel, John T O’Brien, Ph Scheltens, Ian G McKeith, D Williams, C Ballard, and Frederik Barkhof. A comprehensive study of gray matter loss in patients with alzheimer’s disease using optimized voxel-based morphometry. Neuroimage, 18(4):895–907, 2003.

[35] Gary W. Van Hoesen, Josef Parvizi, and Ching-Chiang Chu. Orbitofrontal Cortex Pathology in Alzheimer’s Disease. Cerebral Cortex, 10(3):243–251, 03 2000.

[36] Jeremy D Schmahmann and Deepak N Pandya. The cerebrocerebellar system. International review of neurobiology, 41:31–60, 1997.

[37] Roberta M Kelly and Peter L Strick. Cerebellar loops with motor cortex and prefrontal cortex of a nonhuman primate. Journal of neuroscience, 23(23):8432–8444, 2003.

[38] Jeremy D. Schmahmann. The cerebrocerebellar system: anatomic substrates of the cerebellar contribution to cognition and emotion. International Review of Psychiatry, 13(4):247–260, 2001.

[39] Craig W Berridge and Barry D Waterhouse. The locus coeruleus–noradrenergic system: modulation of behavioral state and state-dependent cognitive processes. Brain research reviews, 42(1):33–84, 2003.

[40] Mara Mather and Carolyn W Harley. The locus coeruleus: essential for maintaining cognitive function and the aging brain. Trends in cognitive sciences, 20(3):214–226, 2016.

[41] Akiko Satoh and Koichi M Iijima. Roles of tau pathology in the locus coeruleus (lc) in age-associated pathophysiology and alzheimer’s disease pathogenesis: potential strategies to protect the lc against aging. Brain research, 1702:17–28, 2019.

[42] Heiko Braak and Kelly Del Tredici. Where, when, and in what form does sporadic alzheimer’s disease begin? Current opinion in neurology, 25(6):708–714, 2012.

[43] Panos Theofilas, Alexander J. Ehrenberg, Sara Dunlop, Ana T. Di Lorenzo Alho, Austin Nguy, Renata Elaine Paraizo Leite, Roberta Diehl Rodriguez, Maria B. Mejia, Claudia K. Suemoto, Renata Eloah De Lucena Ferretti-Rebustini, Livia Polichiso, Camila F. Nascimento, William W. Seeley, Ricardo Nitrini, Carlos Augusto Pasqualucci, Wilson Jacob Filho, Udo Rueb, John Neuhaus, Helmut Heinsen, and Lea T. Grinberg. Locus coeruleus volume and cell population changes during alzheimer’s disease progression: A stereological study in human postmortem brains with potential implication for early-stage biomarker discovery. Alzheimer’s and Dementia, 13(3):236–246, 2017.

[44] Heidi I L Jacobs, David A Hopkins, Helen C Mayrhofer, Emiliano Bruner, Fred W van Leeuwen, Wijnand Raaijmakers, and Jeremy D Schmahmann. The cerebellum in Alzheimer’s disease: evaluating its role in cognitive decline. Brain, 141(1):37–47, 07 2017.

[45] Jinhui Wang, Xinian Zuo, Zhengjia Dai, Mingrui Xia, Zhilian Zhao, Xiaoling Zhao, Jianping Jia, Ying Han, and Yong He. Disrupted functional brain connectome in individuals at risk for alzheimer’s disease. Biological psychiatry, 73(5):472–481, 2013.

[46] Huanqing Yang, Hua Xu, Qingfeng Li, Yan Jin, Weixiong Jiang, Jinghua Wang, Yina Wu, Wei Li, Cece Yang, Xia Li, et al. Study of brain morphology change in alzheimer’s disease and amnestic mild cognitive impairment compared with normal controls. General psychiatry, 32(2), 2019.

[47] Erika J Lampert, Kingshuk Roy Choudhury, Christopher A Hostage, Bharath Rathakrishnan, Michael Weiner, Jeffrey R Petrella, P Murali Doraiswamy, Alzheimer’s Disease Neuroimaging Initiative, et al. Brain atrophy rates in first degree relatives at risk for alzheimer’s. NeuroImage: Clinical, 6:340–346, 2014.

[48] Inbal Maidan, Anat Mirelman, Jeffrey M Hausdorff, Yaakov Stern, and Christian G Habeck. Distinct cortical thickness patterns link disparate cerebral cortex regions to select mobility domains. Scientific reports, 11(1):1– 11, 2021.

[49] Mark A Eckert, Vinod Menon, Adam Walczak, Jayne Ahlstrom, Stewart Denslow, Amy Horwitz, and Judy R Dubno. At the heart of the ventral attention system: the right anterior insula. Human brain mapping, 30(8):2530–2541, 2009.

[50] Xingyun Liu, Xiaodan Chen, Weimin Zheng, Mingrui Xia, Ying Han, Haiqing Song, Kuncheng Li, Yong He, and Zhiqun Wang. Altered functional connectivity of insular subregions in alzheimer’s disease. Frontiers in aging neuroscience, 10:107, 2018.

[51] Megha M Vasavada, Jianli Wang, Paul J Eslinger, David J Gill, Xiaoyu Sun, Prasanna Karunanayaka, and Qing X Yang. Olfactory cortex degeneration in alzheimer’s disease and mild cognitive impairment. Journal of Alzheimer’s disease, 45(3):947–958, 2015.

[52] Mark Rapoport, Robert van Reekum, and Helen Mayberg. The role of the cerebellum in cognition and behavior: a selective review. The Journal of neuropsychiatry and clinical neurosciences, 12(2):193–198, 2000.

[53] William W Seeley, Richard K Crawford, Juan Zhou, Bruce L Miller, and Michael D Greicius. Neurodegenerative diseases target large-scale human brain networks. Neuron, 62(1):42–52, 2009.

[54] Akram Bakkour, John C Morris, David A Wolk, and Bradford C Dickerson. The effects of aging and alzheimer’s disease on cerebral cortical anatomy: specificity and differential relationships with cognition. Neuroimage, 76:332–344, 2013.

[55] Dan D Jobson, Yoshiki Hase, Andrew N Clarkson, and Rajesh N Kalaria. The role of the medial prefrontal cortex in cognition, ageing and dementia. Brain communications, 3(3):fcab125, 2021.

[56] Tao Wang, Feng Shi, Yan Jin, Weixiong Jiang, Dinggang Shen, and Shifu Xiao. Abnormal changes of brain cortical anatomy and the association with plasma microrna107 level in amnestic mild cognitive impairment. Frontiers in Aging Neuroscience, 8:112, 2016.

[57] Y Grignon, C Duyckaerts, M Bennecib, and J-J Hauw. Cytoarchitectonic alterations in the supramarginal gyrus of late onset alzheimer’s disease. Acta neuropathologica, 95(4):395–406, 1998.

[58] Takamasa Yokoi, Hirohisa Watanabe, Hiroshi Yamaguchi, Epifanio Bagarinao, Michihito Masuda, Kazunori Imai, Aya Ogura, Reiko Ohdake, Kazuya Kawabata, Kazuhiro Hara, et al. Involvement of the precuneus/-posterior cingulate cortex is significant for the development of alzheimer’s disease: a pet (thk5351, pib) and resting fmri study. Frontiers in Aging Neuroscience, page 304, 2018.

[59] Marilyn Albert, Yuxin Zhu, Abhay Moghekar, Susumu Mori, Michael I Miller, Anja Soldan, Corinne Pettigrew, Ola Selnes, Shanshan Li, and Mei-Cheng Wang. Predicting progression from normal cognition to mild cognitive impairment for individuals at 5 years. Brain, 141(3):877–887, 01 2018.

[60] Manuela Pietzuch, Anna E. King, David D. Ward, and James C. Vickers. The influence of genetic factors and cognitive reserve on structural and functional resting-state brain networks in aging and alzheimer’s disease. Frontiers in Aging Neuroscience, 11, 2019.

[61] Zhengjia Dai, Qixiang Lin, Tao Li, Xiao Wang, Huishu Yuan, Xin Yu, Yong He, and Huali Wang. Disrupted structural and functional brain networks in alzheimer’s disease. Neurobiology of aging, 75:71–82, 2019.

[62] Farnaz Zamani Esfahlani, Joshua Faskowitz, Jonah Slack, Bratislav Mišić, and Richard F Betzel. Local structure-function relationships in human brain networks across the lifespan. Nature communications, 13(1):1– 16, 2022.

